# Plumbagin-induced oxidative stress leads to inhibition of Na^+^/K^+^-ATPase (NKA) in canine cancer cells

**DOI:** 10.1101/532812

**Authors:** Yousef Alharbi, Arvinder Kapur, Mildred Felder, Lisa Barroilhet, Timothy Stein, Bikash R. Pattnaik, Manish S. Patankar

## Abstract

The Na^+^/K^+^-ATPase (NKA) complex is the master regulator of membrane potential and a target for anti-cancer therapies. Here, we investigate the effect of drug-induced oxidative stress on NKA activity. The natural product, plumbagin increases oxygen radicals through inhibition of oxidative phosphorylation. As a result, plumbagin treatment results in decreased production of ATP and a rapid increase in intracellular oxygen radicals. We show that plumbagin induces apoptosis in canine cancer cells via oxidative stress. We use this model to test the effect of oxidative stress on NKA activity. Using whole-cell patch-clamp electrophysiology we demonstrate that short-term exposure (4 min) to plumbagin results in 48% decrease in outward current at +50 mV. Even when exogenous ATP was supplied to the cells, plumbagin treatment resulted in 46% inhibition outward current through NKA at +50 mV. In contrast, when the canine cancer cells were pre-treated with the oxygen radical scavenger, N-acetylcysteine, the NKA inhibitory activity of plumbagin was abrogated. These experiments demonstrate that the oxidative stress-causing agents such as plumbagin and its analogues, are a novel avenue to regulate NKA activity in tumors.

## 1. Introduction

The Na^+^/K^+^-ATPase (NKA) is a major ion pump that is essential to maintain an optimal membrane potential [1,2] The NKA pumps three sodium ions from inside to the outside of the cell and concurrently transports two potassium ions inside the cell. The transport of both sodium and the potassium ions occurs against their individual concentration gradients. The ATPase activity of the NKA hydrolyses ATP to provide the energy needed for active ion transport.

In cancer cells, there is significant evidence that NKA is expressed at higher levels and the ion transport activity is also enhanced as compared to normal cells [4,5]. There is ample data suggesting that inhibition of NKA activity by cardiac glycosides result in cell death. Therefore, there are initiatives to develop NKA inhibitors as chemotherapeutics for the treatment of cancer. A majority of these studies have focused on digoxin, ouabain and other cardiac glycosides because of their known ability to potently inhibit NKA activity [6–11].

While pre-clinical studies have demonstrated that cardiac glycosides can be used to treat tumors, in the clinical setting it has been found that these agents have higher toxicity when used at concentrations that are required for clinical management of cancer [12,13]. Therefore, new approaches are necessary to target NKA using agents that are potent against the cancer but at the same time have an acceptable safety profile. In the current study, we investigate the natural product, plumbagin, for its NKA inhibitory activity.

Hypoxia results in oxidative stress and is known to damage the NKA complex through at least two mechanisms. The oxidized NKA complex is proteosomally degraded resulting in decrease in the expression of this ion pump on the cell membrane [14–17]. As a result, there is membrane depolarization and hence, cell death. Oxidative stress also is known to activate protein kinase C (PKC) which in turn phosphorylates NKA [14–19]. The phosphorylated form of NKA is internalized from the cell membrane, again resulting in membrane depolarization and cell death (6-9).

These previous observations suggested that NKA complex is vulnerable to hypoxia-induced oxidative stress. We therefore asked if agents that trigger oxidative stress in cancer cells could also negatively influence NKA function. Recently, we have demonstrated that the natural product, plumbagin, increases intracellular oxygen radicals in cancer cells by interfering with mitochondrial electron transport [20]. The oxidative damage caused through treatment with plumbagin results in apoptosis of the cancer cells. While plumbagin is known to affect several different pathways that lead to apoptosis (for example, p53 activation, NFκB and PKCε) we investigated if this molecule can also affect NKA activity because of its ability to initiate an intracellular oxygen radical flux.

Here, we employ canine cancer cells (a model we are developing to obtain preclinical data on the use of plumbagin and its analogs for treatment of solid tumors) to test the effect of plumbagin on NKA activity. Using whole cell patch clamping, we demonstrate that treatment of canine cancer cells with plumbagin results in rapid decrease in NKA activity. Our results confirm that the oxidative stress induced by plumbagin is the reason for the suppression of NKA activity. Based on our results, we propose that when evaluating the chemotoxicity mechanisms of oxidative stress-causing agents such as plumbagin, the loss of NKA activity should also be considered as a contributing mechanism to apoptotic cell death.

## 2. Materials and methods

### 2.1 Antibodies and reagents

Antibodies to caspase 3 and β-actin were purchased from Cell Signaling, CA. Secondary antibodies were from Jackson Immunochemicals, MA. Plumbagin was purchased from Sigma Aldrich, MO. Tissue culture media and other reagents were obtained from ThermoFisher unless stated otherwise.

### 2.2 Cell culture

The canine cancer cell lines, CTAC (Canine thyroid adenocarcinoma) [21], Denny (Canine hemangiosarcoma also referred to as DEN-HAS/Fitz) [3], Payton (Canine Osteosarcoma) [22], and 17CM (Canine Oral Melanoma also known as 17CM98) [23] cells were maintained in DMEM media supplemented with 10% FBS and 100 g/ml streptomycin in humidified incubator at 37°C with 5% CO_2_). The cell lines were incubated with either plumbagin or N-acetylcysteine (NAC) as indicated for each experiment. In all experiments, DMSO was used as the vehicle control.

### 2.3 Cell viability assay

The anti-proliferative activity of plumbagin was assessed by MTT colorimetric assay. Briefly, 10,000 cells were seeded in flat-bottomed 96-well plates and incubated overnight in humidified incubator at 37°C and in 5% CO_2_. The cells were treated with DMSO (vehicle) or with varying concentrations of plumbagin as noted for each experiment. The treatment was conducted for 48 or 72 h. After incubation with the drug or vehicle, the MTT reagent (20 μl of 50 μg/ml stock solution in phosphate buffered saline) was added to each well and the plates were incubated for 3 h at 37°C. DMSO (100 μl) was then added to each well to dissolve the formazan crystals. The plate was mixed gently and the optical density of the solution in each well was determined at 560 nm on a microplate reader.

In some assays, the cells (10,000/well) were incubated with NAC (2 mM) for 30 min. The NAC was removed and each well was washed twice with phosphate buffered saline. After the wash, the cells were treated with vehicle or plumbagin and the MTT assay was conducted as described above.

### 2.4 Western blotting

The canine cancer cells (3-4×10^6^) were plated in 10 cm tissue culture plates. The cells were treated with vehicle (DMSO) or with plumbagin (5-10 μM) for different time points as depicted for each experiment. After treatment, the cells were lysed using RIPA buffer (10 mM Tris-HCl (pH 8.0) containing 1 mM EDTA, 1% Triton X100, 0.1% sodium deoxycholate, 0.1% sodium dodecyl sulfate, 140 mM sodium chloride, 1 mM PMSF and protease inhibitor cocktail (ThermoFisher) and 1 mM sodium orthovanadate).

The cell lysate was sonicated on ice for 25 s and was cleared by centrifugation at 12,000 X g for 30 min. The supernatant was transferred to new tubes and the concentration of the protein was determined by using the BCA kit. Approximately 30 μg of total protein was loaded in each well of the stacking gel and separated by SDS-PAGE. The proteins were transferred to PVDF transfer membrane (1 h transfer at constant current of 250 mA). After transfer, the membranes were blocked with 1XTBST (500 mM Tris base, 150 mM NaCl, 1xTween20, pH7.5) containing 5% powdered milk. The membranes were then incubated overnight with the primary antibodies suspended in 1X TBST buffer and 5% dry milk at 4°C. The membranes were washed three times (10 ml/per 15 min wash) with1X TBST buffer and incubated with HRP-conjugated secondary antibody for 1 h at room temperature. The membranes were washed three times (10 ml/wash) with 1X TBST buffer, overlaid with detection reagent (Femto or Dura kit from ThermoFisher) and the bands were detected using an X-ray film.

### 2.5 Annexin V assay

Apoptosis of the CTAC cells were determine by using FITC-Annexin V Apoptosis Detection kit (BD Pharmingen, San Diego, CA). The cells were treated with 5 μM of plumbagin for 18 h at 37°C. The cells were washed twice with cold phosphate buffered saline and resuspended with 1X binding buffer (10 mM HEPES/NaOH, 140 mM NaCl, 2.5 mM CaCl_2_, pH 7.4). The cells were then transferred to 5 ml tubes, stained with FITC-Annexin V and propidium iodide (PI) and analyzed using flow cytometry on a BD FACSCalibur instrument and the data analyzed using FlowJo software (12).

### 2.6 Intracellular oxygen radical generation

Oxygen radical formation in the canine cancer cell lines in response to plumbagin was monitored as reported previously [20,24,25]. Briefly, cells were labeled with 10□ μM H_2_-DCFDA (Molecular Probes, OR) for 30□ min at 37□ °C, followed by treatment with plumbagin. The cells were washed with PBS, harvested and ROS content was measured on a FACSCalibur (BD Biosciences, CA) flow cytometer. Flow cytometry data were analyzed using FlowJo software (Ashland, OR).

### 2.7 Electrophysiology

CTAC cells were plated on coverslips and placed in incubator at 37°C overnight. For each coverslip, one healthy cell was selected based on its morphology. This healthy cell was patch clamped with a microglass pipet. The pipet measured 5-7 M? when filled with 20 mM tetramethylammonium hydroxide (TMA-OH), 90 mM NaOH, 20 mM tetrathylammonium chloride (TEA-CL), aspartic acid, 2 mM MgCl_2_, 5 mM EGTA, 5 mM Tris-ATP, 2.5 mM Tris-creatine phosphate, 5 mM glucose, 10 mM HEPES at pH 7.4. The bath solution composed of (40 mM NaCl, 5.4 mM KCl, 0.5 mM MgCl_2_, 0.33 mM NaH_2_PO_4_, 5.5 mM glucose, 5 mM HEPES, 2 mM BaCl_2_, 1 mM CsCl, 0.2 mM CdCl_2_, 2 mM NiCl_2_ and 1 μM nifedipine at pH 7.4). This combination of pipette and bath solution allowed us to isolate NKA activity [26]. All recordings were performed in whole-cell patch clamp configuration. Cells were gravity fed (VC-8, Warner Instruments, Hamden, CT) with bath solution until a stable reading for the current was recorded. Once the current was stable, the cell was perfused with 5 µM of plumbagin. The recordings were made using a 2 s ramp protocol from 50 mV to −160 mV applied from a holding potential of –60 mV. All recordings were performed at room temperature using an electrophysiology rig built around a Nikon Eclipse TE300 inverted microscope (Nikon, USA), PATCHSTAR micropositioner (Scientifica, East Sussex, UK), Low noise amplifier Axopatch 200B, D/A converter Digidata 1550B and the data were analyzed using pClamp-10 software (all from Molecular Devices, Sunnyvale, CA).

## 3. Results

### 3.1. Plumbagin inhibits proliferation of canine cancer cells

Recently, we demonstrated that plumbagin is an efficient inhibitor of cancer cell proliferation [27–31]. The naphthoquinone unit of plumbagin likely mimics the quinone ring of ubiquinone (CoQ10) (Fig. 1A) and interferes with electron transport in the oxidative phosphorylation pathway [20]. To obtain support for the use of plumbagin as a chemotherapeutic agent in cancer patients, we are developing pre-clinical data in a canine cancer model. Here, we demonstrate for the first time that similar to our previous studies with human and murine cancer cells, plumbagin also inhibits the proliferation of canine osteosarcoma cell lines. We tested four canine cancer cell lines (17CM, Payton, Denny and CTAC) that were available at our institution. In the MTT assay, we observed that plumbagin inhibited the viability of all four of these cancer cell lines (Fig. 1 B). The data from assays conducted at 48 and 72 h time points demonstrated that plumbagin was significantly potent in decreasing the viability of the cells with an IC_50_ between 5-10 µM (Fig. 1 B).

**Fig. 1.**
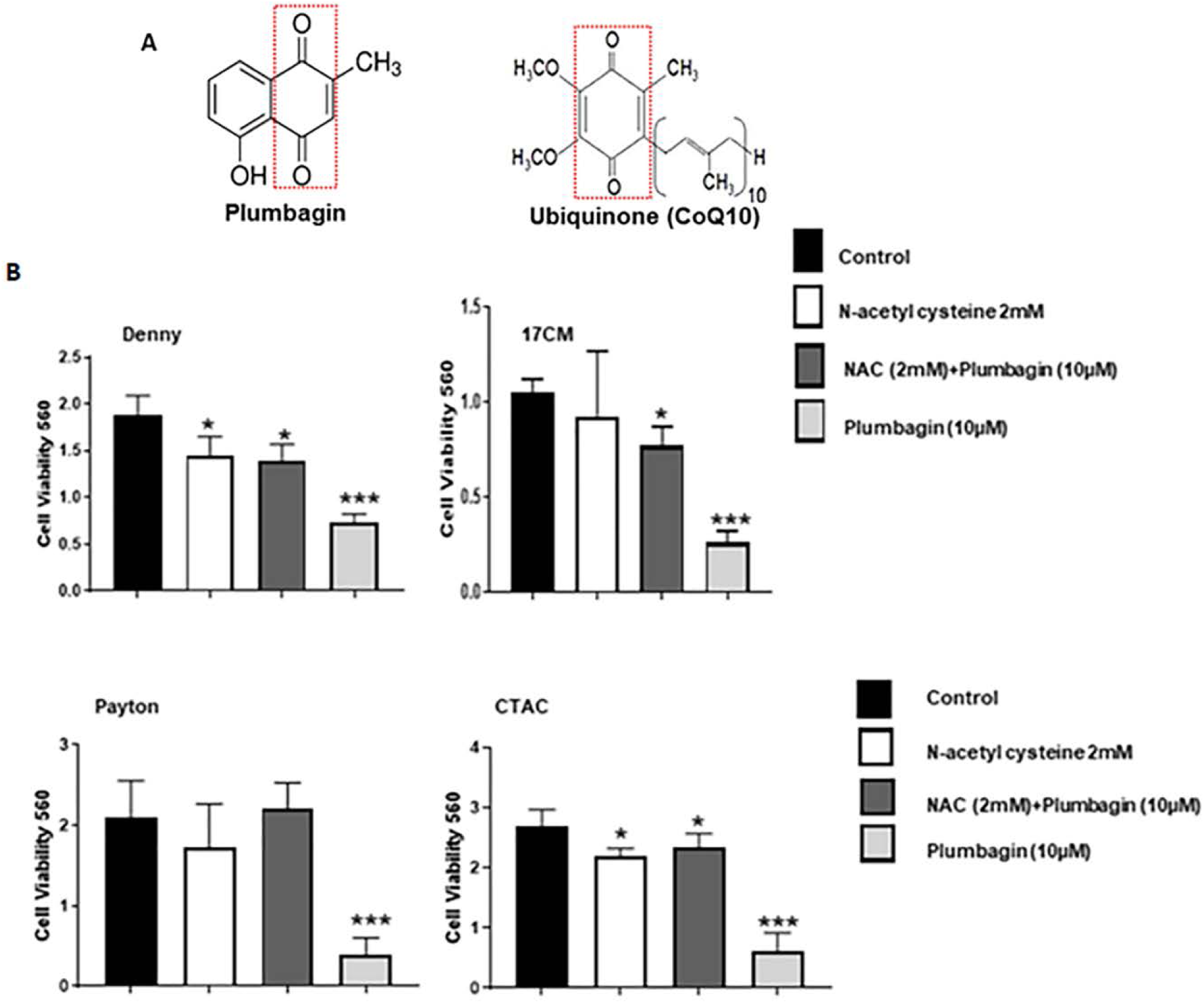
Plumbagin inhibits the proliferation of canine cancer cells. (**A**) Structural similarities between plumbagin and ubiquinone are indicated by the red box. (**B**) The canine cancer cell lines (CTAC, 17CM, Denny and Peyton) were treated with plumbagin (0–10 μM) for 48 and 72 hours and proliferation of the cell lines was determined using MTT assay. The average of eight wells was used to plot the bar graphs for each condition tested. Data shown is representative of three independent biological replicates. * p= 0.013, ** p=0.003, and *** p=0.0002 of test versus controls for each time point.

### 3.2. Plumbagin treatment causes cell death by apoptosis

Next, we investigated if the decrease in proliferation of plumbagin-treated cancer cells was due to cell death via apoptosis. The CTAC cells were treated with plumbagin (5 µM) overnight. After treatment, the cells were harvested and stained with Annexin V-FITC and propidium iodide. Flow cytometry of the cells demonstrated significant increase in Annexin V-positive population, indicating cell death was occurring via apoptosis (Fig. 2 A, B). Apoptosis in CTAC, Denny and Payton cells upon exposure to plumbagin (5 µM for 24, 48 and 72 h) was confirmed by the increase in expression of cleaved caspase 3 detected by western blotting (Fig. 2 C).

**Fig. 2.**
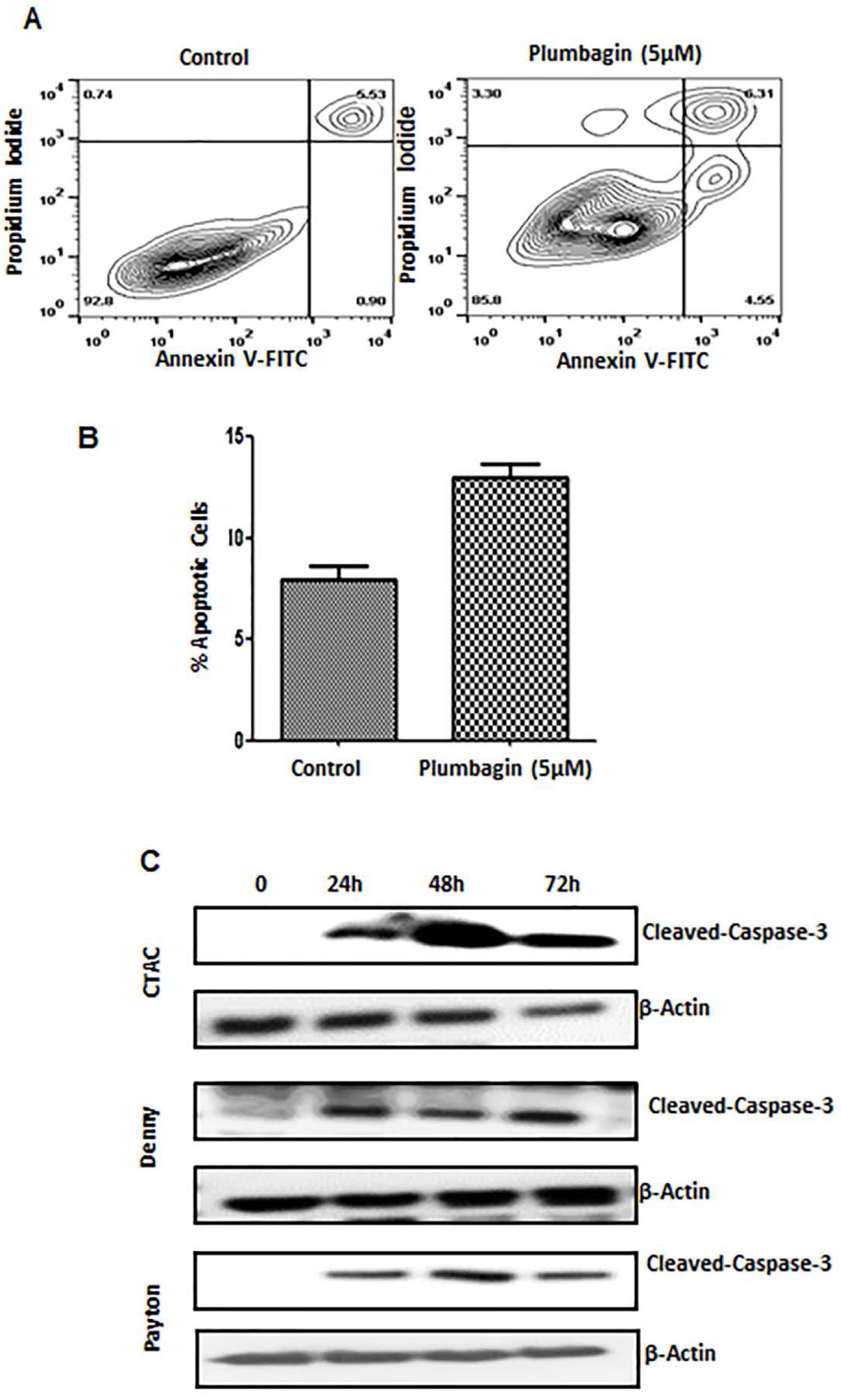
Plumbagin induces apoptosis in canine cancer cells. **A)** CTAC cells were treated with plumbagin (5 μM) for 24 h. Following treatment, the cells were stained with Annexin V-FITC (to detect expression of phosphoserine on the surface of the cell) and propidium iodide (as a live/dead cell indicator) and the cells were analyzed by analytical flow cytometry. **B)** The bar chart shows average data from three independent replicates for the annexin V-FITC/propidium iodide assays. (p= 0.006). **C)** Three canine cancer cell lines CTAC, Denny and Payton were treated with plumbagin (5 μM) for the depicted time points. After treatment, the cells were lysed and expression of cleaved caspase 3 and β-actin (as a loading control) was determined by western blotting. The blots shown are representative of three independent replicates.

### 3.3. Plumbagin causes oxidative stress in canine cancer cells

Our experiments with human cancer cell lines showed that plumbagin induces oxidative stress [20]. When the CTAC cells were labeled with the ROS sensing fluorescent dye, H2DCFDA and treated with plumbagin (5 µM), a rapid increase in intracellular oxygen radicals was detected by flow cytometry analysis of the cells (Fig. 3 A, B). Time-dependent monitoring of the cells by flow cytometry indicated a rapid rise in intracellular ROS at 15 min of treatment, the first timepoint tested. Higher levels of intracellular ROS were maintained till 2 h of treatment, the last timepoint tested in our assays (Fig. 3 A, B).

**Fig. 3.**
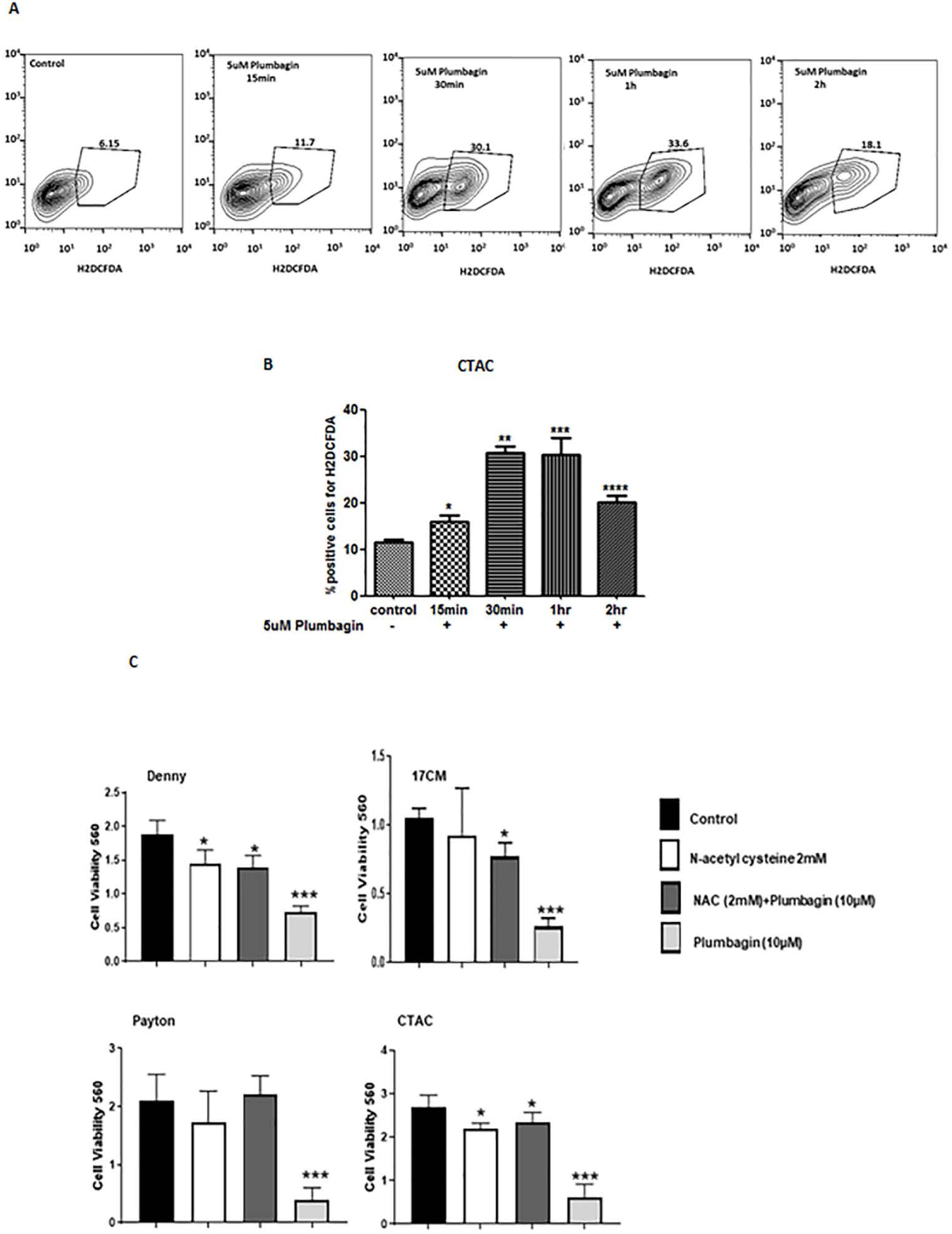
Plumbagin induces oxidative stress in canine cancer cells. **A)** CTAC cells were pre-labeled with the oxygen radical sensing dye, H2DCFDA. The cells were then exposed to plumbagin (5 μM) for the depicted time points. The release of oxygen radicals in the cells was quantified by analytical flow cytometry. * p= 0.04, ** p=0.0001, *** p=0.002 and **** p=0.001 of test versus controls for each time point. **B)** Bar graphs show average data from three independent replicate experiments. The percent of cells showing increase in oxygen radicals was calculated from the flow cytometry datasets. **C)** MTT assays were conducted to demonstrate that neutralization of the intracellular oxygen radicals by pre-loading the cells with 2 mM N-acetylcysteine (NAC) for 30 min attenuated plumbagin activity. Cells were washed to remove excess NAC from media and then treated with plumbagin (5 μM) or vehicle for 48 h. MTT assays were conducted to determine effect of the treatments on proliferation of each cell line. Data shown is representative of three independent replicates. * p=0.003 and *** p=0.0002 of test versus controls for each time point.

### 3.4. Plumbagin-induced oxidative stress is responsible for decreased proliferation

N-Acetylcysteine (NAC) is a potent oxygen scavenger and a substrate for glutathione synthesis [32–35]. We therefore pretreated cells with NAC for 30 min. The cells were washed to remove excess NAC from the media prior to adding plumbagin. This step was undertaken to ensure that NAC in the media could not pre-react and chemically inactivate plumbagin before its transport to the cancer cells. Under these conditions, the anti-proliferative effect of plumbagin was significantly reduced as indicated by the increase in cell viability in MTT assays that were conducted with the four canine cancer cells, Denny, Payton, 17CM and CTAC (Fig. 3 C).

### 3.5. Plumbagin inhibits NKA

After demonstrating that plumbagin was inducing cell death in the canine cancer cell lines, we next focused our attention to studying the effect of plumbagin on NKA activity. To test this directly, we conducted whole cell patch clamping experiments on CTAC cells in presence of pipette and bath solution that isolated NKA current.

When the CTAC cells were exposed to plumbagin (10 µM), the time course showed a decrease in both inward and outward current within 20 minutes of drug application (Fig. 4A). The outward current at +50 mV in the untreated cells (Fig. 4 A, black trace) was 1835 pA whereas upon plumbagin treatment (Fig. 4A, red trace), the current decreased to 780 pA. Similarly, the inward current at −150 mV in the untreated CTAC (Fig. 4A, black trace) and plumbagin-treated (Fig. 4A, red trace) was 795 pA and 619 pA, respectively.

**Fig. 4.**
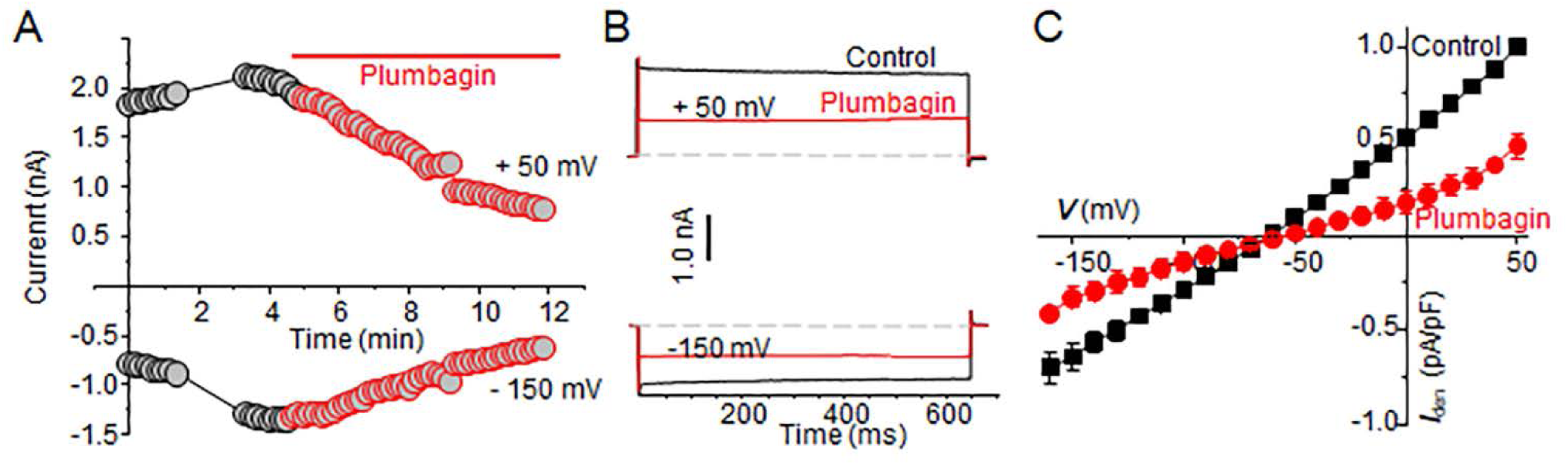
Plumbagin inhibits NKA channel currents. **A)** The time course measurement of current amplitude in a representative CTAC cell before and after plumbagin (5 μM) treatment. For these recordings, current was measured at +50 mV (as outward current) and −150 mV (as inward current) from a holding potential of −60 mV. **B)** Representative current trace during control (black traces) and plumbagin (5 µM) treated (red traces) CTAC cell is shown at either +50 mV or −150 mV from a holding potential of −60 mV. Dotted lines represent zero current. **C)** The average of 3 cells of normalized current-voltage (I-V) curve in control (black trace) and after plumbagin (5µM, Red trace) application are shown (n = 3).

To determine if the current was time or voltage dependent, we applied step-wise voltage gradient. As shown in Fig. 4 B, neither outward nor inward current was time or voltage dependent between −150 to +50 mV. The current at +50 mV in the control experiments was 1835 pA whereas in the plumbagin-treated CTAC cells, the current decreased to 780 pA (Fig. 4 B). When the voltage was maintained constant at −150 mV, the current measured was −795 pA. In comparison, plumbagin treatment decreased the inward current measured at a constant voltage of −150 mV to 619 pA (Fig. 4 B). These results indicated that plumbagin was inhibiting NKA-mediated ion transport. Average current voltage plots from three recorded cells demonstrate the inhibitory activity of plumbagin on NKA (Fig. 4 C). These average current voltage (I-V) curves show inhibition of outward current at +50 mV by about ∼ 52.3%.

### 3.6. Inhibition of NKA by plumbagin is not due to decrease in intracellular ATP

We recently demonstrated that plumbagin decreased the intracellular ATP levels in cancer cells [20]. We therefore tested if the inhibition of NKA observed in plumbagin-treated CTAC was due to decreased availability of intracellular ATP. To address this possibility, we conducted electrophysiology experiments where the pipet solution was supplemented with 5 mM ATP. This ensured that there was sufficient intracellular ATP for optimal NKA activity.

Even upon ATP supplementation, plumbagin continued to inhibit the inward and outward currents in CTAC cells (Fig. 5A-C). The outward current at +50 mV in the untreated cells measured 1925 pA and upon treatment with plumbagin the current decreased to ∼807 pA (Fig. 5A, back trace). Similarly, at −150 mV, the current measured in untreated CTAC was 1273 pA whereas in the plumbagin treated cell the current was decreased to 519 pA (Fig. 5A, red trace). This inhibition of the NKA activity by plumbagin in the presence of ATP supplementation was also observed when the voltage was applied in a stepwise gradient (Fig. 5B). The average of five cells plotted in current-voltage curve (I-V), showed that plumbagin inhibits NKA activity by ∼58.3%.

**Fig. 5.**
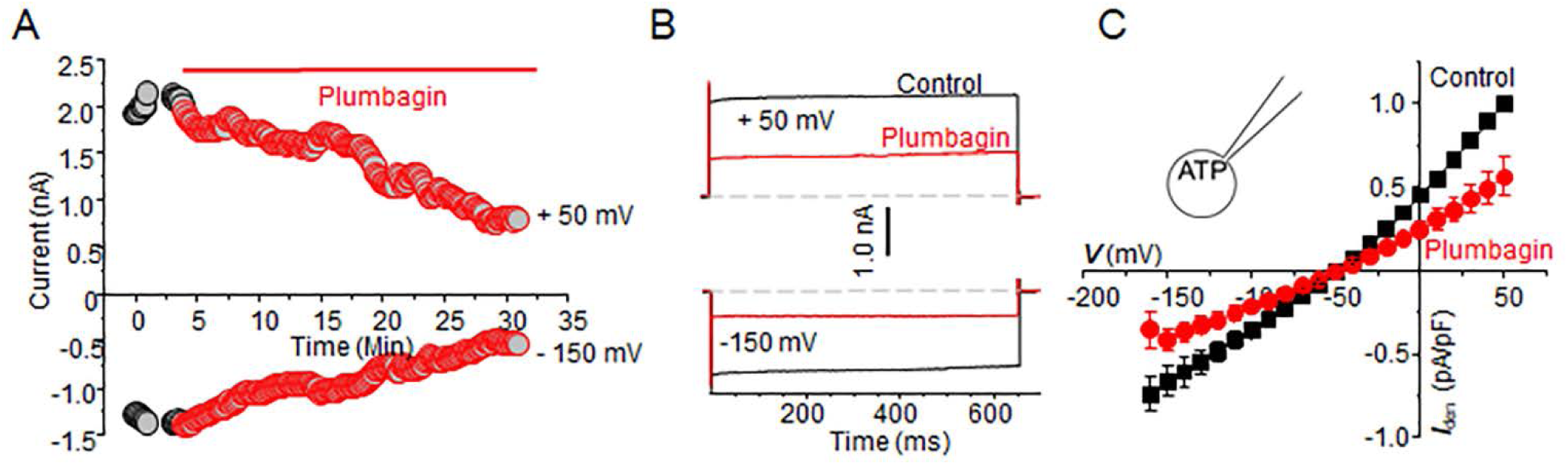
ATP did not prevent NKA inhibition by plumbagin. **A)** The time course plot for the control and plumbagin (5 μM) treated CTAC cell is shown using ATP (5 mM) supplemented pipette solution. **B)** Voltage steps plot at +50 and –160 mV shows inhibition of NKA activity even in the presence of ATP in the pipet solution (trace color same as in 4B). **C)** The normalized current-voltage plot for all three, control (black trace) and plumbagin-treated (red trace) CTAC cells are shown. The application of the voltage steps and current measurement are identical to those used in Fig. 4.

### 3.7. NKA inhibitory activity is abrogated by oxygen radical scavenger, N-acetylcysteine

The NKA complex is sensitive to reactive oxygen radicals (6-11). We therefore reasoned that the inhibition of NKA activity was a downstream effect of plumbagin, occurring due to the damage of this ion transport complex by the intracellular oxygen radical flux.

CTAC cells were pre-loaded for 1 h with the oxygen radical scavenger, NAC (1 mM). After washing away the excess NAC, the cells were patched and subsequently treated with vehicle (control) or 10 µM plumbagin. The pretreatment with NAC resulted in almost complete loss of NKA inhibition by plumbagin (Fig. 6 A, B). Loss of the inhibitory activity of plumbagin was also observed in experiments where the cells were subjected to a stepwise voltage gradient (Fig. 6 B). The average of five cells plotted in current-voltage (I-V) curves showed no inhibition of NKA activity by plumbagin in cells pretreated with NAC.

**Fig. 6.**
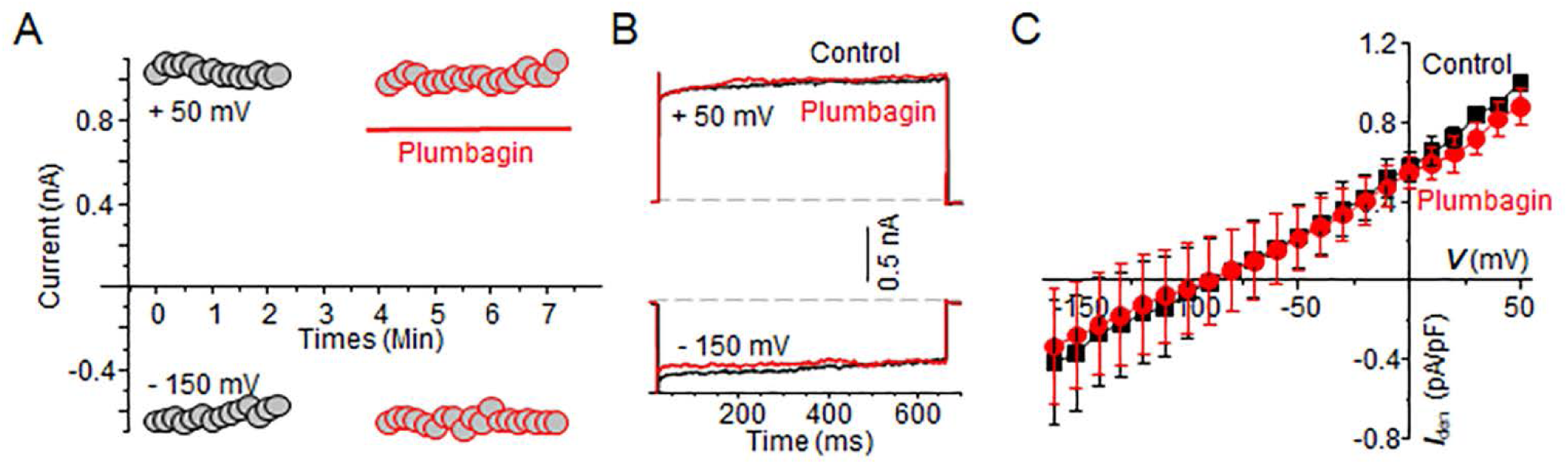
Oxidative stress caused by plumbagin inhibits NKA activity. CTAC cells were preincubated with 2 mM of NAC for 1 hour. Healthy cells were patch clamped before and after treatment with plumbagin (5 μM). **A)** Time course plot of the control and plumbagin treated cell is shown measured at +50 and –150 mV. **B)** Current measured in response to voltage steps of +50 and −150 mV is shown. **C)** Average current-voltage plots (n = 5) for the control (black trace) and plumbagin (5 μM) treated (red trace) CTAC cells are shown. The voltage pulses applied and color scheme are identical to those described in (Fig. 4).

Comprehensive analysis of the electrophysiology experiments clearly demonstrated that while intracellular ATP concentrations had no effect, inhibition of oxidative stress abrogated the NKA-suppressive activity of plumbagin (Fig. 7). The inhibition of the outward current measured at +50 mV after treatment with plumbagin was 47.7 ± 6.6 % (p 0.000866) as compared to control cells (Fig. 7). When exogenous ATP was supplied through the pipet solution, plumbagin continued to significantly inhibit the outward current at +50 mV (46.4 ± 4 % inhibition; p= 0.000177) as compared to control (Fig. 7). In comparison, when the CTAC cells were pre-treated with NAC and then exposed to plumbagin, the outward current measured at +50 mV was comparable to that recorded in the control cells (87.4 ± 9.3 %; p= 0.306605) (Fig.7).

**Fig. 7.**
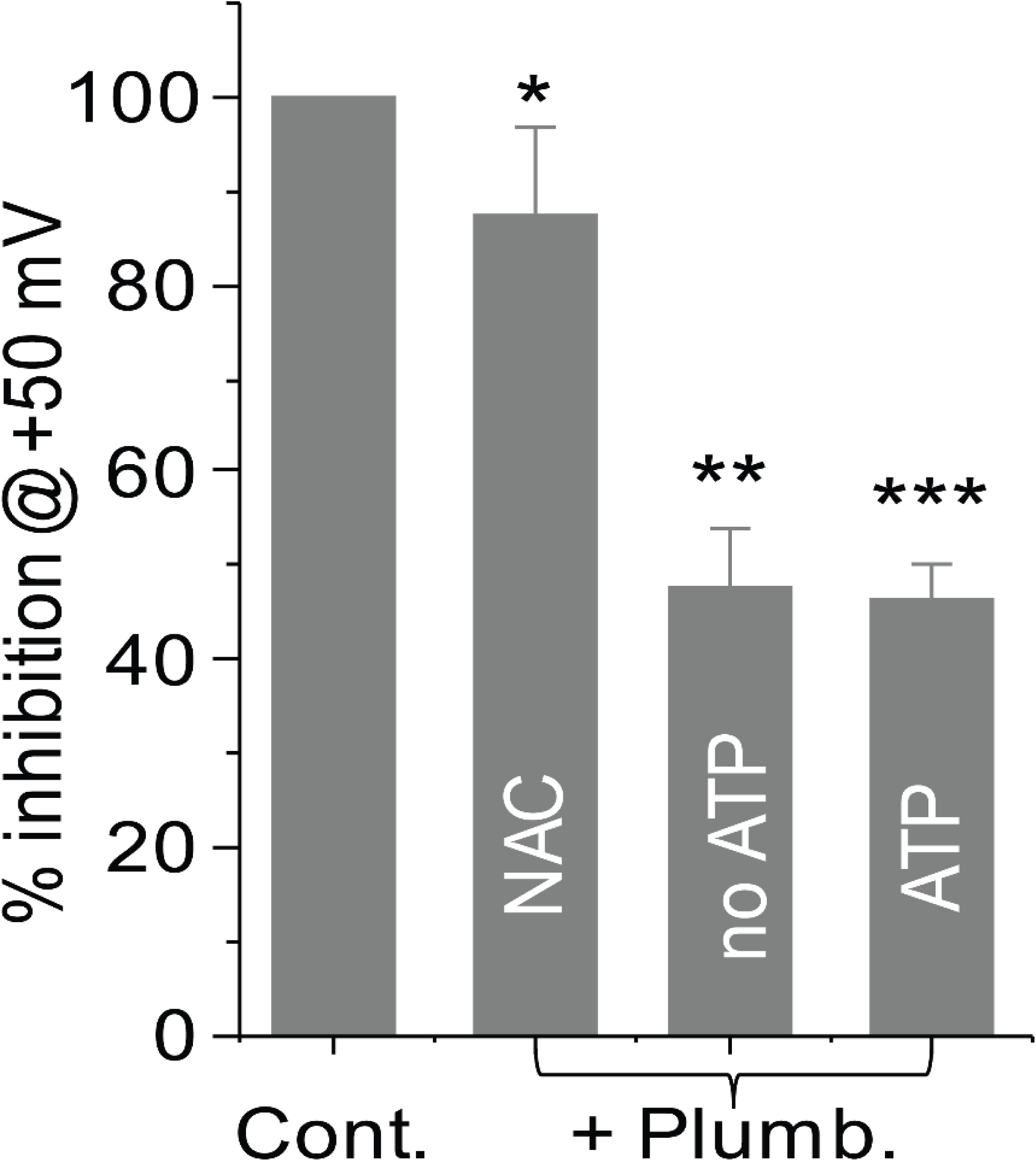
Oxidative stress is responsible for NKA inhibitory activity of plumbagin. The average percent inhibition of the outward current measured at +50 mV in CTAC cells treated with vehicle, plumbagin (5 µM), plumbagin (5 µM with ATP (5 mM-supplied in pipet solution) and cells pretreated with NAC (2 mM) prior to exposure with plumbagin (5 µM) is plotted. Each bar is average data obtained from > 3 individual CTAC cells. * p= 0.306605, ** p=0.000866 and *** p= 0.000177 compared to vehicle control.

## 4. Discussion

Plumbagin is a potent anti-cancer agent that causes cell death in prostate, breast, ovarian and colon cancer cells [20,27,36–42]. In the current study, we demonstrate for the first time that plumbagin can also induce apoptotic cell death in canine cancer cell lines. These results indicate that plumbagin can be further explored in canine cancer models to test for its therapeutic activity in dogs. The preclinical data obtained from such canine cancer models can be used to support the clinical use of plumbagin for the treatment of human cancers.

In addition to demonstrating the anti-cancer effects of plumbagin in the canine model, we also demonstrate that plumbagin activity resulted in the inhibition of NKA. Within minutes of treatment of cells with plumbagin (4 min as tested in our current study), there occurs a significant inhibition of NKA pump activity. Both the inward current (transport of potassium ions into the cell) and outward current (transport of sodium ions from inside to the outside of the cell) is inhibited during this relatively short-term treatment of the cancer cells with plumbagin. To the best of our knowledge, the inhibitory effect of plumbagin on ion transport through the NKA has not been previously described.

Previous studies with plumbagin have indicated that this molecule produces its anti-cancer effects by targeting several important cell signaling pathways. For example, in our studies we demonstrated that plumbagin activates the tumor suppressor, p53. Others have shown that cell death caused by plumbagin is through the inhibition of PKCε, PI3K, pAKT, pJAK-2 and pSTAT3 [29,36,37]. Plumbagin also prevents the transcription factors AP-1, NFκB and Stat3 from interacting with the DNA [43,44]. Downstream effects of plumbagin include inhibition of the anti-apoptotic protein, BCL-XL (13). Clearly, these studies show that plumbagin affects several different pathways that can contribute to apoptosis. Our results demonstrate that inhibition of NKA should be added to the list of potential molecules that are altered by plumbagin.

Just as our studies demonstrate that oxidative stress induced by plumbagin is responsible for inhibition of NKA activity, previously we have also shown that the increase in oxygen radicals is required for activation of p53. Whether oxidative stress in plumbagin-treated cells also leads to inhibition of PKCε, NFκB and other molecules remains to be determined.

Novel therapies are being designed to target NKA in tumors [45,46]. Maintaining an optimum electrical potential is essential for healthy cells. The higher reliance of cancer on NKA activity creates an opportunity for development of anti-cancer therapies. To date the majority of the approaches investigated to inhibit NKA are with molecules that compete for the ion binding site located in the extracellular domain of this ion pump. Our studies with plumbagin are indicating an alternate approach to targeting NKA. This approach involves the use of agents or strategies that will lead to an increase in intracellular oxidative stress. Plumbagin can serve as a prototype molecule for further development of agents that can target NKA activity through such an indirect mechanism of action.

Our proposed approach makes use of the fact that NKA is sensitive to oxidative stress-either through direct damage caused to the protein complex or via phosphorylation by PKC [16,17]. The net effect is that NKA expression is significantly downregulated in cells that are expressing oxidative stress [16,17]. The decreased expression of NKA translates to an inability of the cell to maintain its membrane potential [47]. Since several of the biochemical reactions and pathways are required for the maintenance of an optimal membrane potential, the cell with an improper NKA activity is susceptible to cell death [10,48]. While other ion channels present in the cell membrane can be activated to compensate and maintain ion balance, the deficiency of NKA cannot be completely overcome as it alters equilibrium for both K^+^ and Na^+^. In other words, decreasing NKA activity through oxidative damage results in a cellular insult that is difficult to repair.

In the current study, we have used a direct assay to measure the activity of NKA in the control and plumbagin-treated cells. While this assay clearly indicates that NKA activity is attenuated upon plumbagin treatment, we have not directly shown that this loss of activity is the result of decreased expression of NKA on the canine cancer cells. Antibodies that efficiently recognize NKA subunits from the canine species will be needed for such experiments to monitor expression of the subunits of this ion pump.

The inhibition of NKA activity by plumbagin also creates opportunities for new combination therapies with cardiac glycosides, the classical NKA activity inhibitors. The cardiac glycosides have high potency and therefore have a narrow therapeutic window. We are in the process of developing analogues of plumbagin that can be specifically targeted to tumors. Such agents can be used to selectively generate oxidative stress in cancer cells and thereby attenuate NKA activity. Since the NKA activity will already be compromised by such agents through the induction of oxidative stress, it will create a window of opportunity to follow-up with lower concentrations of cardiac glycosides to further target the residual activity of the NKA and cause even more potent apoptosis in the cancer cells. Our on-going studies are exploring such novel combination approaches as small molecule strategies against ovarian and other solid tumors.

## Acknowledgements

We are deeply grateful to Dagna Sheerar (University of Wisconsin-Madison Paul P. Carbone Comprehensive Cancer Center, Flow Cytometry Facility) for advice and assistance with the flow cytometry assays. YA was supported by Saudi Arabia Government scholarship and by the Ride fellowship award from the University of Wisconsin-Madison. Studies were supported by funds from the Department of Obstetrics and Gynecology and the Wisconsin Ovarian Cancer Alliance to MSP and LB and R01EY024995 and MD Matthews Research Professorship from Retina Research Foundation to BRP. Core grant (CA14520) support to the University of Wisconsin Comprehensive Cancer Centers Flow Cytometry facility is acknowledged.

## Author’s contributions

Yousef Alharbi: Designed and conducted the experiments and co-wrote the manuscript. Arvinder Kapur: Conducted the flow cytometry experiments. Mildred Felder: Assisted with cell culture. Lisa Barroilhet: Provided intellectual input. Tim Stein: Developed the canine cancer cells. Bikash Pattnaik: Supervised the electrophysiology experiments and helped in designing the experiments, analysis and writing the manuscript. Manish Patankar: Designing the experiments, interpreting data and writing of the manuscript.

